# NMPylation and de-NMPylation of SARS-CoV-2 Nsp9 by the NiRAN domain

**DOI:** 10.1101/2021.06.13.448258

**Authors:** Bing Wang, Dmitri Svetlov, Irina Artsimovitch

## Abstract

Nsp12, the catalytic subunit of SARS-CoV-2 RNA-dependent RNA polymerase (RdRp), contains two active sites that catalyze nucleotidyl-monophosphate (NMP) transfer (NMPylation). RNA synthesis is mediated by the RdRp active site that is conserved among all RNA viruses and has been a focus of mechanistic studies and drug discovery. The second active site resides in a Nidovirus RdRp-Associated Nucleotidyl transferase (NiRAN) domain. Both catalytic reactions are essential for viral replication, but the mechanism and targets of NiRAN are poorly characterized. One recent study showed that NiRAN transfers NMP to the first residue of RNA-binding protein Nsp9. Another study reported a structure of SARS-CoV-2 replicase with an extended Nsp9 in the NiRAN active site but observed NMP transfer to RNA instead. We show that SARS-CoV-2 Nsp12 efficiently and reversibly NMPylates the native but not the extended Nsp9. Substitutions of the invariant NiRAN residues abolish NMPylation, whereas a substitution of a catalytic RdRp Asp residue does not. NMPylation is inhibited by nucleotide analogs, pyrophosphate, and bisphosphonates, suggesting a path for rational design of NiRAN inhibitors. We hypothesize that Nsp9 remodels both active sites of Nsp12 to support initiation of RNA synthesis by RdRp and subsequent capping of the product RNA by the NiRAN domain.

## INTRODUCTION

Coronaviruses (CoVs) are single-stranded positive-sense (+) RNA viruses that constitute the *Coronaviridae* family in the order *Nidovirales* (1). CoVs cause many respiratory and gastrointestinal infections in humans, from mild common colds to severe respiratory diseases, including the ongoing COVID-19 pandemic (2,3). Severe acute respiratory syndrome coronavirus 2 (SARS-CoV-2), the etiological agent of COVID-19, is the third zoonotic CoV to have caused a major disease outbreak in humans in the last two decades (4). The prevalence of CoVs in animal reservoirs argues that future viral pandemics are all but certain (3,5) and makes advance preparations imperative. While the availability of effective vaccines against SARS-CoV-2 has been a game changer for the current COVID-19 pandemic, broad-spectrum antiviral drugs are needed to protect unvaccinated and immunocompromised individuals and to buy time needed for the development of new vaccines at the onset of the next viral epidemic.

CoVs have very large (approximately 30 kb) genomes that encode non-structural proteins (Nsps) required for viral gene expression and replication. Upon infecting human cells, the SARS-CoV-2 RNA genome is translated to produce a long polyprotein that is cleaved into Nsps 1 through 16 by the viral protease Nsp5 (6). Nsp12, the catalytic subunit of RNA-dependent RNA polymerase (RdRp) stands out as the only protein that is present in all RNA viruses (7) and is therefore an attractive target for broad-spectrum antivirals (8). The high degree of conservation of the RdRp structure and its key catalytic elements (7,9) encouraged efforts to repurpose existing antivirals, such as remdesivir (10,11) and favipiravir (12), for the treatment of COVID-19 (13). However, these pursuits have not yet produced an effective clinical treatment, suggesting that a detailed mechanistic analysis of the viral replication cycle may be required to identify the best points for intervention.

SARS-CoV-2 RdRp is composed of the catalytic Nsp12 and accessory Nsp7 and Nsp8 proteins, forming a stable four-subunit Nsp12•7•8_2_ holoenzyme (14–16). The RdRp holoenzyme associates with the superfamily 1 helicase Nsp13 (17) to form a minimal Nsp12•7•8_2_•13_2_ replication-transcription complex (RTC). Other proteins, including a proofreading exonuclease Nsp14, capping enzymes, and RNA-binding proteins Nsp9 and Nucleocapsid are thought to associate with the RTC in a large multi-subunit complex that mediates synthesis and modification of all viral RNAs (6). Nsp12 also contains a second catalytic module, an N-terminal 250-residue <u>Ni</u>dovirus <u>R</u>dRp-<u>A</u>ssociated <u>N</u>ucleotidyl transferase (NiRAN) domain (9). The NiRAN domain displays significant sequence divergence as compared to RdRp, with only four conserved motifs (preA_N_, A_N_, B_N_, and C_N_) comprising the NiRAN signature (18). The NiRAN domain is present in all nidoviruses but has no homologs in other RNA viruses and, together with the helicase (HELD) domain, is a genetic marker for the order *Nidovirales* (18).

As first shown with an equine arteritis virus (EAV) enzyme from the *Arteriviridae* family of nidoviruses, RdRp self-NMPylates *in vitro* with a clear preference for UTP as a substrate and the Mn^2+^ ion as a cofactor (18). Substitutions of several conserved NiRAN residues abolish nucleotidyl transfer *in vitro* and abrogate EAV replication in cell culture to the same extent as do substitutions of the catalytic RdRp residues (18). These results demonstrate that the NiRAN domain plays a critical role in the viral life cycle and thus is a valid target of antiviral drug discovery. Subsequent studies of two viruses from *Coronaviridae*, HCoV-229E and SARS-CoV-2, led to similar conclusions (19).

The location of the NiRAN nucleotidyl transfer site is well established, but the identity of the NMP acceptor remains debated. Single-particle cryogenic electron microscopy (cryoEM) studies of SARS-CoV-2 RdRp (17,20) revealed nucleotides bound to Nsp12 residues shown to be required for self-NMPylation *in vitro* (18). A finding that NiRAN active site is structurally homologous to that of a protein pseudokinase, selenoprotein O/SelO (16,17,21), supported the proposed role of NiRAN in covalent NMPylation of protein targets (18). Consistently, NiRAN-dependent NMPylation of the N-terminal primary amine of the Asn1 residue of SARS-CoV-2 Nsp9 has been demonstrated *in vitro* (19); this modification was proposed to underpin *de novo* initiation of RNA synthesis *in vivo* and is essential for viral replication (19). Furthermore, in a cryoEM structure of Nsp9 bound to the SARS-CoV-2 RTC (20), the Nsp9 Asn1 is adjacent to the NiRAN active site (**Fig. 1A**). Strikingly, however, this study failed to detect Nsp9 modification; instead, Nsp12 transferred GMP to the 5′ terminus of an RNA oligonucleotide, a reaction proposed to initiate the capping pathway of the viral mRNA (20). Nsp9 inhibited this nucleotidyl transfer reaction and was proposed to guide the modified RNA for further processing (20).

**Figure 1.**
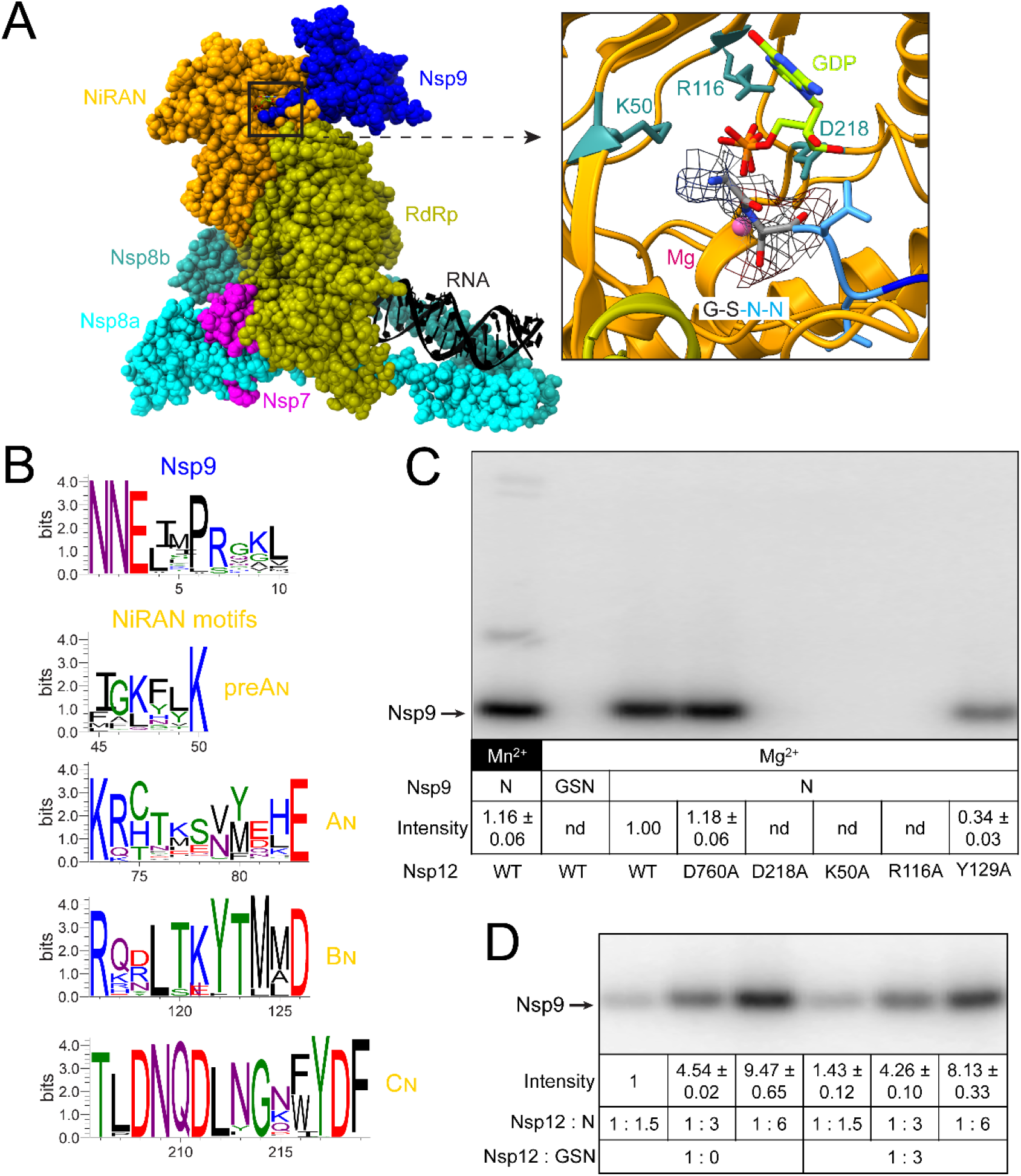
NiRAN-mediated NMPylation of Nsp9. **A**. Cryo-EM structure of the SARS-CoV-2 RTC bound to inert Nsp9 with a two-residue extension at the N-terminus (20). Left. Overall structure of the complex; PDB: 7CYQ; Nsp13 helicase is not shown. Proteins are shown as molecular surfaces and RNA as cartoon. The color coding corresponds to the figures throughout this manuscript unless otherwise specified. Right. Zoom in on the active site of the NiRAN domain (AS2) with GDP-Mg^2+^ (lime carbon atoms and magenta sphere, respectively). Side chains of key conserved residues from the pre-A_N_, B_N_, and C_N_ motifs that were substituted in this work are shown as sticks. Four N-terminal residues of Nsp9 (GSNN) are shown; the cryo-EM difference density for Gly and Ser residues from EMDB: 30504 is shown (gray mesh). Structural figures were prepared with Coot (57), UCSF ChimeraX 1.2, and PyMOL Molecular Graphics System, version 2.4.1, Schrodinger, LLC. **B**. Conservation of residues at the N-terminus of Nsp9 and four conserved NiRAN motifs in alpha-, beta-, gamma-, and deltacoronavirus genera. **C**. Mutations in the NiRAN active site and Nsp9 N-terminal GS extension abolish NMP transfer, but Mn^2+^ is dispensable. NMPylation efficiency was compared to that observed with the wild-type Nsp12 in the presence of 1 mM Mg^2+^ (set at 1) and is shown as mean ± SD (n=3); nd, no signal detected above the background. **D**. Multi-round NMPylation of Nsp9 is not inhibited by the inert ^GSN^Nsp9. NMPylation efficiency was compared to that observed with ^N^Nsp9 present at 1.5 x molar excess over Nsp12 in the absence of ^GSN^Nsp9 (set at 1) and is shown as mean ± SD (n=3).

Our findings that noncognate NTPs and nucleoside analogs modulate RdRp activity suggested that the RdRp and NiRAN active sites (thereafter referred to as AS1 and AS2, respectively) could be allosterically connected (22). Testing this hypothesis necessitates parallel assays of both nucleotidyl transfer activities and in turn requires using cognate NiRAN substrates. In agreement with HCoV-229E studies by Ziebuhr and colleagues (19), we show that SARS-CoV-2 Nsp12 efficiently NMPylates Nsp9 that has the native N-terminus, but not an Nsp9 variant that bears two additional N-terminal residues. Substitutions of the invariant NiRAN residues abolished Nsp9 NMPylation, whereas substitution of a catalytic RdRp residue, Asp760, did not. We found that this reaction proceeds equally efficiently with Mg^2+^ and Mn^2+^, is largely insensitive to the identity of the NTP, and is inhibited by nucleotides, pyrophosphate and bisphosphonates. The latter findings suggest a starting point for identification of NiRAN inhibitors.

## MATERIALS AND METHODS

### Construction of expression vectors

Plasmids used in this study are shown in **Table S1**. The SARS-CoV-2 *nsp7/8/9/12* genes were codon-optimized for expression in *E. coli* and synthesized by GenScript and subcloned into standard pET-derived expression vectors under control of the T7 gene 10 promoter and *lac* repressor. The derivative plasmids were constructed by standard molecular biology approaches with restriction and modification enzymes from New England Biolabs. DNA oligonucleotides or vector construction and sequencing were obtained from Millipore Sigma, synthetic DNA fragments for Gibson Assembly – from IDT. The sequences of all plasmids were confirmed by Sanger sequencing at the Genomics Shared Resource Facility (The Ohio State University). All plasmids are deposited to Addgene.

### Protein expression and purification

The expression and purification of Nsp7/8/12 and Nsp12 mutants are described in our previous study (22). All purification steps were carried out at 4 °C. Nsp9 variants were overexpressed in *E. coli* BL21 (DE3) cells (Novagen, Cat#69450). Cells were grown in lysogenic broth (LB) with kanamycin (50 μg/mL). Cells were cultured at 37 °C to an OD_600_ of 0.6-0.8 and the temperature was lowered to 16°C. Expression was induced with 0.1 mM isopropyl-1-thio-β-D-galactopyranoside (IPTG; Goldbio, Cat#I2481C25) for 18 hours. Induced cells were harvested by centrifugation (6000 × *g*), resuspended in lysis buffer (50 mM HEPES, pH 7.5, 300 mM NaCl, 5 % glycerol (v/v), 1 mM Phenylmethylsulfonyl fluoride (PMSF; ACROS Organics, Cas#329-98-6), 5 mM β-ME, 10 mM imidazole), and lysed by sonication. The lysate was cleared by centrifugation (10,000 × *g*). The soluble protein was purified by absorption to Ni^2+^-NTA resin (Cytiva, Cat#17531801), washed with Ni-buffer A (50 mM HEPES, pH 7.5, 300 mM NaCl, 5 % glycerol, 5 mM β-ME, 50 mM imidazole), and eluted with Ni-buffer B (50 mM HEPES, pH 7.5, 300 mM NaCl, 5 % glycerol, 5 mM β-ME, 500 mM imidazole). The eluted protein was further loaded onto a desalting column (Cytiva, Cat#17508701) in desalting buffer (50 mM HEPES, pH 7.5, 5 % glycerol, 5 mM β-ME, 300 mM NaCl) to remove imidazole. The fusion protein was treated with TEV protease (for pIA1364) or SUMO protease (for pIA1414) at 4 °C overnight. The sample was supplemented with 20 mM imidazole and passed through Ni^2+^-NTA resin. The untagged protein was loaded onto a Sephacryl S-100 HR column (Cytiva, Cat#17116501) in desalting buffer. Peak fractions were assessed by SDS–PAGE and Coomassie staining. Purified protein was dialyzed into storage buffer (20 mM HEPES, pH 7.5, 150 mM NaCl, 45 % glycerol, 5 mM β-ME), aliquoted, and stored at −80°C.

### Conservation analysis

A total of 75 viral genomes, representing alpha-, beta-, gamma-, and deltacoronaviruses, were fetched from Uniprot reference Proteomes (version 2021_02) (23). Multiple sequence alignment (MSA) was done by MAFFT (version 7) (24). WebLogo (version 3) (25) was used to generate the sequence logos based on the MSA.

### Nsp9 NMPylation

All NMPylation assays were carried out at 37 °C. For standard NMPylation assay, 0.5 μM Nsp12 and 5 μM Nsp9 were incubated in NMPylation buffer (25 mM HEPES, pH 7.5, 15 mM KCl, 5 % glycerol, 2 mM MgCl_2_, 2 mM DTT) for 5 min, then 25 μM GTP and 10 μCi [α^32^P]-GTP (PerkinElmer, Cat#BLU006H250UC) were added to start the reaction. After a further 10 min incubation, the reaction was mixed with 4X NuPAGE™ LDS Sample Buffer (ThermoFisher, Cat#NP0007).

### Competition assays

To assess the competition of ^GSN^Nsp9 with ^N^Nsp9, Nsp12 was incubated with ^GSN^Nsp9 for 5 min. Then ^N^Nsp9 (at desired concentrations), 25 μM GTP, and 10 μCi [α^32^P]-GTP were added to start the reaction. For nucleotide competition assays, NTPs, NTP analogs, and pyrophosphate were used at 0.5 mM. After incubation of Nsp12/9 for 5 min in NMPylation buffer, the competitor NTPs (Cytiva, Cat#27202501), GDP (Sigma-Aldrich, Cat#G7127), GMP (Sigma-Aldrich, Cat#G8377), Inosine-5′-Triphosphate (ITP; TriLink Biotechnologies, Cat#N-1020), ppGpp (Trilink Biotechnologies, Cat#N-6001), GpCpp (Jena Bioscience, Cat#NU-405S), Remdesivir triphosphate (RTP; MedChemExpress, Cat#GS443902), and pyrophosphate (PP_i_; Sigma-Aldrich, Cat#71515) was added together with 25 μM GTP and 10 μCi [α^32^P]-GTP to start the reaction. After a further 10 min incubation, the reactions were stopped with LDS Sample Buffer as above.

### De-NMPylation

To determine if PP_i_ can reverse the NMPylation reaction, 0.5 μM Nsp12, 5 μM Nsp9, 25 μM GTP and 10 μCi [α^32^P]-GTP were incubated in NMPylation buffer for 15 min, then 0.5 mM PP_i_ was added. To determine which active site of Nsp12 is responsible for the de-NMPylation activity, 0.5 μM His-tagged Nsp12, 20 μM Nsp9, 50 μM GTP, and 10 μCi [α^32^P]-GTP were incubated in NMPylation buffer (2 mM DTT in the buffer was replaced by 2 mM β-mercaptoethanol) for 20 min. Dynabeads (ThermoFisher, Cat#10103D) were added to remove the His-tagged Nsp12, followed by adding 0.5 μM Nsp12 variants and 0.5 mM PP_i_. Samples were quenched at indicated time points and analyzed by electrophoresis.

### Inhibition by bisphosphonates

0.5 μM Nsp12 and 5 μM Nsp9 were incubated with different concentrations of Risedronate (Sigma-Aldrich, Cat#PHR1888) or Foscarnet (Sigma-Aldrich, Cat#PHR1436) in the NMPylation buffer for 5 min, then 25 μM GTP and 10 μCi [α^32^P]-GTP were added to start the reaction. Reactions were performed for 10 min.

### RNA extension and cleavage

An RNA oligonucleotide (5′ -UUUUCAUGCUACGCGUAGUUUUCUACGCG-3′; 4N) with Cyanine 5.5 at the 5′-end was obtained from Millipore Sigma (USA). The RNA scaffold was annealed in 20 mM HEPES, pH 7.5, 50 mM KCl by heating to 75 °C and then gradually cooling to 4 °C. To test RdRp activity, reactions were carried out at 37 °C with 500 nM Nsp12 variants, 1 μM Nsp7, 1.5 μM Nsp8, 250 nM RNA, and 250 μM NTPs in the transcription buffer (20 mM HEPES, pH 7.5, 15 mM KCl, 5 % glycerol, 1 mM MgCl_2_, 2 mM DTT) for 20 min at 37 °C. For pyrophosphorolysis, holo RdRp was preincubated the RNA scaffold at 37 °C for 5 min in the transcription buffer; then the indicated combinations of PP_i_ and NTPs were added. Reactions were stopped by adding 2 × stop buffer (8 M Urea, 20 mM EDTA, 1X TBE, 0.2 % bromophenol blue).

### Sample analysis

Protein samples were heated for 5 min at 95 °C and separated by electrophoresis in NuPAGE™ 4-12 % gels (ThermoFisher, Cat# NP0329BOX). RNA samples were heated for 2.5 min at 95 °C and separated by electrophoresis in denaturing 9 % acrylamide (19:1) gels (7 M Urea, 0.5X TBE). The gels were visualized and quantified using Typhoon FLA9000 (GE Healthcare) and ImageQuant. All assays were carried out in triplicates. The means and standard deviation (SD) were calculated by Excel (Microsoft).

## RESULTS

### NMPylation requires the native N-terminus of SARS-CoV-2 Nsp9

The NIRAN domains are very divergent, apart from four short signature motifs (18). Consistent with a hypothesis that protein NMPylation is a cognate activity of the NiRAN domain, Mn^2+^-dependent self-NMPylation of RdRps from EAV, HCoV-229E and SARS-CoV-2 viruses has been reported (18,19). Such an activity is common among AMPylases, which frequently transfer AMP to their autoinhibitory domains (26). Substitutions of invariant NiRAN residues abolished self-NMPylation in these RdRps (18,19), confirming the NiRAN role therein and prompting a search for other NiRAN targets.

Ziebuhr and colleagues recently showed that HCoV-229E and SARS-CoV-2 Nsp12s efficiently transfer NMPs to Nsp9 (19), a small (113 residues) RNA-binding protein that is essential for viral replication (27–29). Nsp9 and Nsp12 modifications shared the requirements for NTP substrates, metal cofactors, and NiRAN residues, arguing that both reactions utilize similar mechanisms. Mass spectrometry identified the primary amine of the N-terminal Asn, which is conserved among CoVs (**Fig. 1B**), as a site of Nsp9 modification (19). Mutational analysis revealed that (*i*) the Asn2 residue was critical for modification; (*ii*) Asn1 could be substituted with Ala or Ser with a modest loss of reactivity; and (*iii*) the presence of even one additional N-terminal Ala residue abolished Nsp9 NMPylation (19). In support of the essential role of its NMPylation, Nsp9 substitutions had parallel effects on NMP transfer *in vitro* and on viral replication (19).

In a cryoEM structure of SARS-CoV-2 RTC bound to Nsp9, Asn1 residue is positioned near the NIRAN-bound GDP•BeF_3_^−^ (**Fig. 1A**) and Asn2, which is essential for modification of HCoV-229E Nsp9 (19), makes contacts to residues in the NiRAN and palm domains (20). However, Yan *et al*. did not detect Nsp9 modification and instead observed GMP transfer to RNA, which they proposed represents a key early step in the capping pathway (20). The lack of Nsp9 reactivity is most likely explained by the presence of two additional, non-native residues, Gly and Ser, at the N-terminus of the recombinant Nsp9 used to obtain the structure. While these residues were not modeled in PDB:7CYQ, the GSNNELSPVALR tryptic peptide was identified by mass-spectrometry analysis and the density for Gly-2/Ser-1 residues is discernible in the EM map (**Fig. 1A**). In the presence of these additional residues, the cognate NMPylation site, the N1 amine, is eliminated. Different metal ion cofactors, protein tags, or other reaction variables could also explain discrepancies in observed Nsp12 catalytic properties.

Ideally, one would want to assay both nucleotidyl transfer activities under identical conditions. In our experiments, we used standard solution conditions that support efficient RNA synthesis (**Fig. S1**), [α^32^P]-GTP, which supports efficient Nsp9 modification (18,19) as an NMP donor; and SARS-CoV-2 Nsps containing native N- and C-termini.

We first assayed NMP transfer of Nsp12 alone using two Nsp9 proteins: a variant with the native N-terminus (^N^Nsp9; confirmed by MS analysis) and Nsp9 with two additional residues at −1 and −2 (^GSN^Nsp9), identical to that used in (20). These recombinant proteins were produced by cleavage of tagged Nsp9 precursors by Ubiquitin-like-specific protease 1 (Ulp1) and Tobacco Etch Virus (TEV) proteases, respectively. We observed efficient GMP transfer to ^N^Nsp9 but not to ^GSN^Nsp9 by the wild-type (WT) Nsp12 (**Fig. 1C**). NMPylation was abrogated by substitutions of conserved NiRAN residues (K50A in preA_N_, R116A in B_N_, and D218A in C_N_; **Fig. 1B**) that inhibit viral replication in cell culture (19), but not by the D760A substitution in AS1. As expected, NiRAN mutants did not affect RNA synthesis, whereas the D760A variant was inactive (**Fig. S1**). The Y129A substitution at the NiRAN/RdRp domain interface modestly reduced both activities (**Figs. 1C** and **S1**). We conclude that, as shown for the HCoV-229E system (19), SARS-CoV-2 Nsp9 that has the native N terminus is efficiently NMPylated by AS2.

Our findings and those of Slanina *et al*. (19) underscore the potential importance of native termini to protein function. The N- and C-termini are commonly modified to include purification tags, an approach that is justified when these ends are phylogenetically variable. However, in the context of CoV protein maturation, the “correct” ends are generated upon proteolytic cleavage of the polyproteins by the viral protease. Given that the very first residue of Nsp9 is the target of NMPylation, the potential importance of the identity of this residue is obvious, as is the prudence of preserving its native identity in experiments unless and until that identity is proven to be unimportant.

### NMPylation occurs in the presence of Nsp7/8 cofactors and does not require Mn^2+^

To match the previously published conditions, we carried GMPylation assays with Nsp12 alone. To ascertain that this activity is preserved in context of the transcribing RdRp holoenzyme (Nsp12•7•8_2_), we repeated our assays in the presence of Nsp7, Nsp8, and an RNA scaffold under conditions that support robust RNA extension by SARS-CoV-RdRp (9,22). Our results demonstrate comparable Nsp9 modification by Nsp12 alone or as part of an active transcription complex (**Fig. S2**).

Unlike that of its structural homolog SelO (21), the NiRAN domain’s activity was thought to be dependent on the Mn^2+^ ion, at least for the EAV and HCoV-229E RdRps (18,19). Surprisingly, we observed equally efficient GMPylation in the presence of 1 mM Mg^2+^ or 1 mM Mn^2+^ (**Fig. 1C**), concentrations that correspond to physiological levels of Mg^2+^ but exceed those of Mn^2+^ (30). Mg^2+^ is the major cellular cofactor in electrophilic catalysis, in part due to its superior bioavailability and environmental abundance (30). Although Mn^2+^ can also function as the cofactor for the nucleotidyl transfer reaction for diverse nucleic acid polymerases, Mn^2+^ binding alters the active site geometry (30) to promote base misincorporation (31) and other inefficient reactions (32). The Mn^2+^ ion overrides a requirement for a canonical signal to fortuitously activate cyclic GMP-AMP [cGAMP] synthase (33) and can resuscitate a catalytically-compromised RNA polymerase II (34). Although Mn^2+^ and Mg^2+^ can support similar octahedral coordination in the active site (30), Mn^2+^ has also been observed to form a strikingly different network, independent of the catalytic triad residues (33). Consistent with the Mn^2+^-induced gain-of-function, we observed GMP transfer to BSA in the presence of 1 mM Mn^2+^ (**Fig. S3**). These transfer reactions are very inefficient when compared to NMPylation of Nsp9 but could be easily mistaken for the “cognate” modification when observed in the absence of a real target; we used a 15-min exposure to phosphor screen to obtain an image used in Figure 1.

We do not know why only the Mn^2+^-dependent NMP transfer was observed with EAV and HCoV-229E RdRps since we used very similar reaction conditions (18,19). We speculate that differences in RdRp folding could explain Mn^2+^ dependence. CoV RdRps are highly dynamic enzymes that undergo large conformational changes during the transcription cycle (35) and can become misfolded during expression in heterologous hosts (22). In particular, the NiRAN domain has been captured in different conformational states in cryoEM structures and becomes more ordered upon ligand binding to the active site (9,14,17,36,37).

### Nsp9 modification is not required for its release from Nsp12

Enzymes that mediate protein NMPylation frequently have low affinity for their targets, necessitating covalent linkage of enzyme:substrate complexes for structural analysis (38). The modified Nsp9 appears to be readily released from RdRp: increasing the SARS-CoV-2 Nsp9:12 ratio leads to increased GMP transfer (**Fig. 1C**), and multi-round NMPylation of HCoV-229E Nsp9 has also been reported (19). The formation of a stable complex between SARS-CoV-2 RTC and the modification-resistant ^GSN^Nsp9 (20) raises a possibility that NMPylation is a prerequisite for Nsp9 release.

To test this idea, we used competition between the native ^N^Nsp9 and the inert ^GSN^Nsp9 variant (**Fig. 1C**). We found that preincubation of Nsp12 with a three-fold molar excess of ^GSN^Nsp9 only slightly inhibited modification of ^N^Nsp9 (**Fig. 1D**). Although it is possible that NMPylation alters Nsp9 affinity for RdRp, a possibility that we intend to evaluate in the future, we conclude that free Nsp9 is in a dynamic equilibrium with the Nsp9•12 complex regardless of the presence of the N-terminal nucleotide moiety.

Although the precise role of Nsp9 modification in the viral life cycle remains to be elucidated, the essentiality of Nsp9 modification for viral replication (19) makes NMPylation a valid target for inhibition. Our results indicate that peptidomimetic compounds that resemble the Nsp9 N-terminus are unlikely to serve as efficient inhibitors of NMPylation. However, substrate analogs that bind to AS2 may either interfere with NMP transfer to Nsp9 or lead to modified but non-functional Nsp9.

### SARS-CoV-2 NiRAN can bind diverse nucleotides

Studies of NMPylation have revealed differences in substrate utilization among RdRps. EAV RdRp displayed a strong preference for UTP, followed by GTP, whereas ATP and CTP were barely used (18). By contrast, HCoV-229E RdRp utilized all NTPs with preference for UTP (19). Structures of SARS-CoV-2 transcription complexes with NiRAN-bound nucleotides do not reveal any base-specific contacts (**Fig. 2A**), suggesting that all NTPs would be used as substrates for NMPylation. We tested this assumption using competition experiments in which [α^32^P]-GMP transfer to Nsp9 was assayed in the presence of cold NTPs (**Fig. 2B**). Our results show that while GTP and UTP are marginally more effective competitors, the differences among all NTPs are small, a finding that is at odds with the published preference of EAV and HCoV-229E RdRp for UTP. It is possible that sequence divergence of NiRAN NTP-binding sites could explain their differing substrate preferences; for example, the His75 residue in the A_N_ motif, which contacts ADP•AlF_3_ in the SARS-CoV-2 RTC structure (17), is represented by Cys in HCoV-229E and by Val in EAV NiRAN domains. Structures of nucleotide-bound EAV and HCoV-229E that could help to address this question are not yet available. It is also possible that the assay design contributes to the observed discrepancies. Commercial radiolabeled NTP preparations contain impurities that compromise some sensitive assays, in contrast to high-purity NTPs (see Methods) that we use for all *in vitro* transcription experiments. Using competition of highly purified NTPs against the same radiolabeled NTP substrate minimizes concerns about variable purity of four different [α^32^P]-NTPs and also reduces the cost.

**Figure 2.**
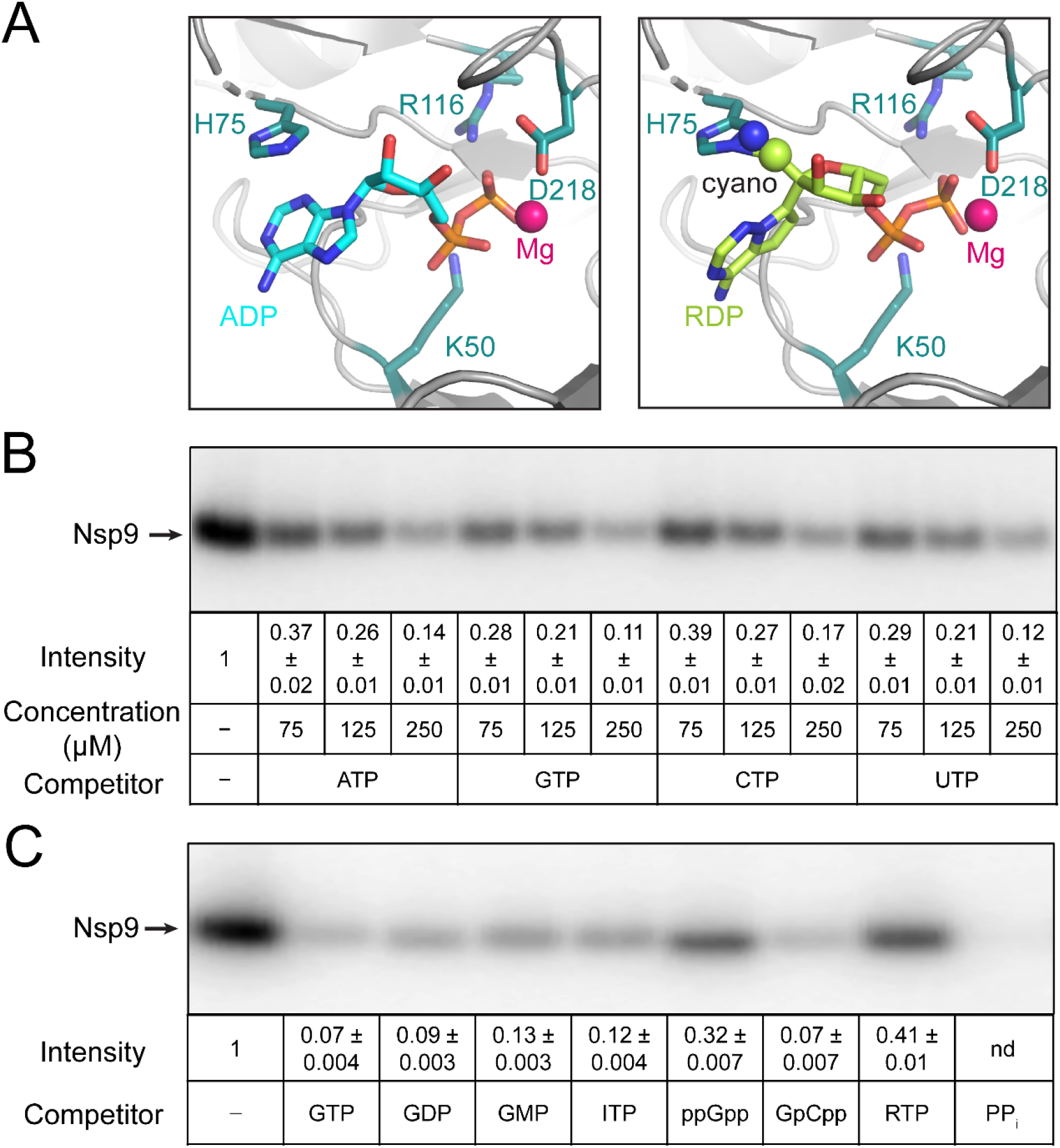
Effects of nucleotides on Nsp9 modification. **A**. Nucleotide binding to the NiRAN active site. The Mg^2+^ ion is shown as magenta sphere and key NiRAN residues - as sticks. Left, ADP (cyan carbon atoms) from PDB:6XEZ. Right, RDP (lime carbon atoms) modeled in place of ADP; the C1’ cyano group of RDP could clash with the side chain of Nsp12 His75. **B**. Unlabeled natural NTPs compete with [α^32^P]-GTP for transfer to Nsp9. **C**. GTP analogs and PP_i_ inhibit NMPylation, but ppGpp and RTP do so less effectively; all nucleotides were present at 0.5 mM. In B and C, NMPylation efficiency was compared to that observed in the absence of competitors, set at 1, and is shown as mean ± SD (n=3).

These results suggest that cellular Nsp9 modification will be controlled by the relative abundance of natural NTPs and that other nucleotides could serve as substrates or competitors of the NMPylation reaction. While some nucleotides could act solely as competitive inhibitors, others (e.g., the ATP analog remdesivir triphosphate; RTP) could transfer the NMP moiety to Nsp9, with a potential to interfere with yet-to-be determined function of Nsp9 in viral replication. To test this idea, we used several nucleotide analogs as competitors of Nsp9 GMPylation. We found that GDP, GMP, ITP (inosine triphosphate), and GMPCPP efficiently competed with [α^32^P]-GMP transfer to Nsp9, whereas ppGpp was less effective (**Fig. 2C**). Surprisingly, and in contrast to ATP (**Fig. 2B**), we found that RTP was a poor competitor (**Fig. 2C**). Analysis of RNA synthesis by SARS-CoV-2 RdRp demonstrated that RTP binds the RdRp active site with much higher affinity than ATP and is a better substrate than ATP (39). Why does RTP fail to compete with GTP during NMPylation? In RTP, a cyano-group is attached to the 1’ position of the ATP ribose sugar; while the cyano-group does not interfere with RMP incorporation into the nascent RNA, it clashes with the Ser861 residue in Nsp12 after RdRp adds three more nucleotides downstream of RMP, leading to a temporary stall during RNA chain extension (11,40). When remdesivir diphosphate is modeled in place of ADP into the structure of SARS-CoV-2 RTC with the NiRAN-bound ADP•AlF_3_ (**Fig. 2A**, right), the cyano-group at the 1’ position clashes with His75, potentially explaining why RTP is a poor substrate for, and competitor of, the NMPylation reaction. Unexpectedly, we also observed that the NMPylation reaction was strongly inhibited when inorganic pyrophosphate PP_i_ was present along with the GTP substrate (**Fig. 2C**, last lane).

### Reversal of NMPylation by pyrophosphate

The nucleotidyl transfer reactions of AS1 and AS2 generate two products: PP_i_, in each case, and either an NMP-adduct (a one-nucleotide-extended RNA) or NMP-Nsp9, respectively. A reverse reaction, pyrophosphorolysis, is unfavorable at physiological concentrations of NTPs and PP_i_, but is commonly used to evaluate the translocation register of multi-subunit DNA-dependent RNA polymerases (41), and has also been observed in RdRps (42,43). RNA polymerases behave as thermal ratchets that oscillate between the pre- and post-translocated registers on the template (44). This motion is rectified by binding of the incoming substrate NTP, which binds in the acceptor site (**Fig. 3A**) and locks the post-translocated state, or of PP_i_, which induces cleavage of the 3′-terminal nucleotide in the product site when the enzyme is in the pre-translocated register (**Fig. 3A**). PP_i_ cleavage leads to shortening of the nascent RNA by one nucleotide and subsequent backward translocation, sometimes in several successive steps (45). The nascent RNA cleavage typically requires superphysiological concentrations of PP_i_ because the transcription elongation complex is biased toward the post-translocated state at most template positions, for bacterial RNA polymerases and SARS-CoV-2 RdRp alike (35,44). Consistently, we observed that scaffold-assembled SARS-CoV-2 complexes were relatively resistant to pyrophosphorolysis even in the absence of NTPs (**Fig. 3A**), a result that is comparable to those obtained with hepatitis C virus (HCV) RdRp (42,43). Interestingly, we observed non-canonical PP_i_-induced RNA cleavage by two nucleotides in a fraction of complexes, reminiscent of reverse pyrophosphorolysis by noncognate NTP substrates in HCV RdRp that also generates a 2-nt cleavage product (42). Similar to the results obtained for the HCV enzyme (42), when PP_i_ was present in 200-fold molar excess over NTPs, polymerization reaction was favored and no cleavage was apparent (**Fig. 3A**); unlike HCV RdRp, SARS-CoV-2 RdRp did not cleave the nascent RNA in the presence of ATP, which induced reverse pyrophosphorolysis most efficiently with HCV RdRp (**Fig. S4**).

**Figure 3.**
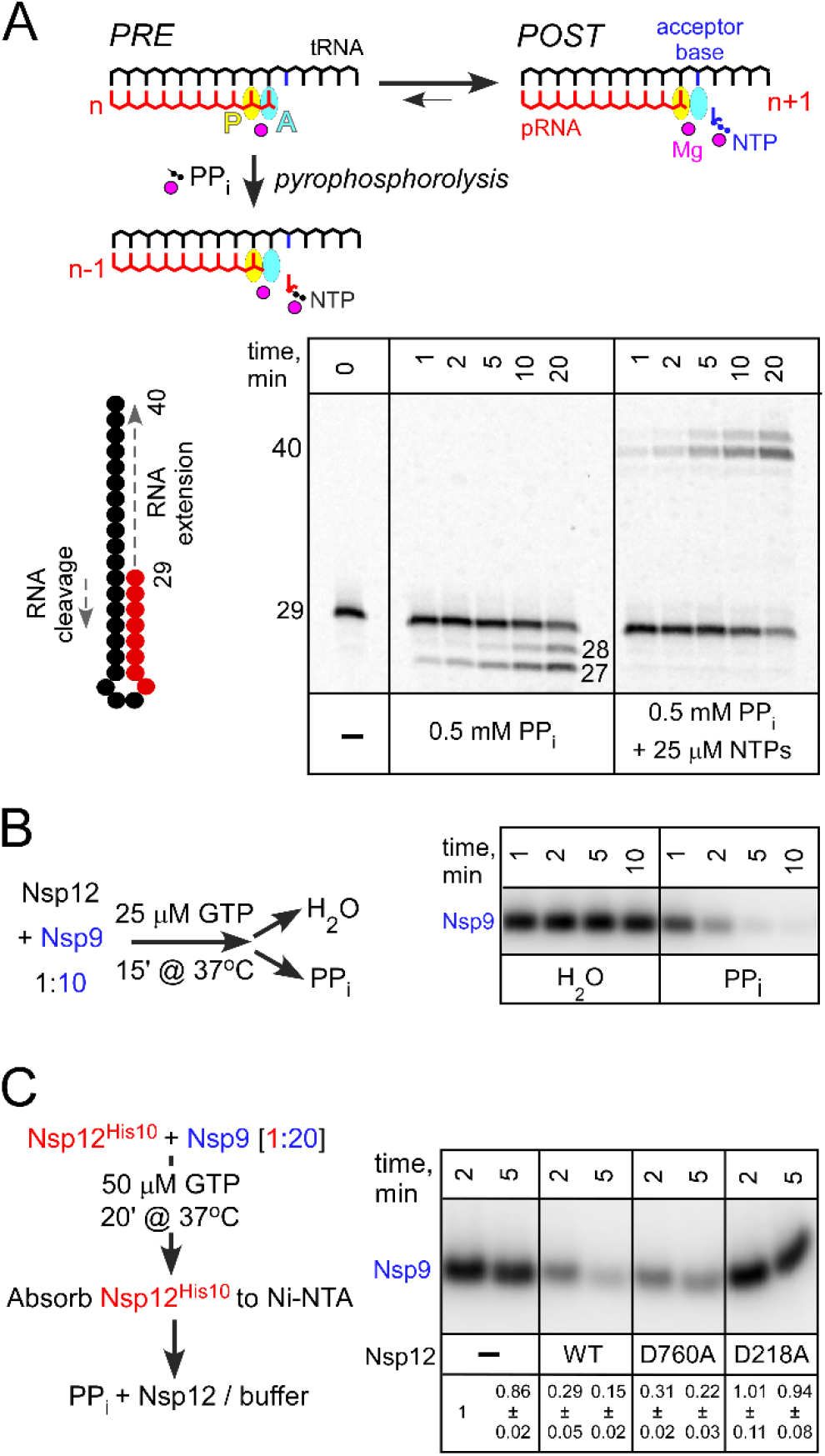
Reversal of NMP transfer reactions in the presence of pyrophosphate. **A**. Top: Pyrophosphorolysis of the nascent product RNA (pRNA; red). The active site (marked by the position of the catalytic Mg^2+^;magenta sphere) consists of two sub-sites: the P-site (product; yellow) and the A-site (acceptor; cyan). Following nucleotide addition, the pRNA 3′ end is bound in the A-site in the pre-translocated state. Upon forward translocation, the 3′ end moves to the P-site and the incoming substrate NTP (blue) can bind to the A-site through base pairing with the acceptor base of the RNA template strand. The pre-translocated state is sensitive to pyrophosphorolysis, which generates the NTP product and a one-nt shortened pRNA. Bottom: SARS-CoV-2 RdRp transcription complex assembled on the hairpin template is completely resistant to PP_i_ even in the presence of NTPs. **B**. Nsp9 NMPylation is reversible by PP_i_. **C**. The hand-over assay in which pre-GMPylated Nsp9 is incubated with WT or mutant Nsp12 variants. Signal intensity was compared to that observed with Nsp9 incubated with buffer (set at 1) and is shown as mean ± SD (n=3).

By contrast, we observed that PP_i_ efficiently inhibited Nsp9 NMPylation even in the presence of the substrate GTP (**Fig. 2C**). This result suggests that PP_i_ binds to AS2, as observed in SARS-CoV-2 RdRp/favipiravir complex (36), with higher affinity than to AS1. Given that Nsp9 binding to the Nsp12 is very dynamic (**Fig. 1C**), PP_i_ could be expected to reverse Nsp9 NMPylation. To evaluate this possibility, we tested if PP_i_ can de-NMPylate ^32^P-GMP-Nsp9. We preincubated Nsp9 with Nsp12 under multi-round reaction conditions prior to addition of PP_i_ (or water). In the presence of 0.5 mM PP_i_, we observed rapid disappearance of the labeled Nsp9 (**Fig. 3B**), indicating that NMPylation is reversible.

In Nsp12, two active sites mediate NMP transfer. A model in which NMPylated Nsp9 serves as a primer for RNA synthesis implies that Nsp9 binds to AS1 and positions the NMP for extension (19). Thus, both active sites could in principle mediate the PP_i_-driven de-NMPylation. To evaluate the contribution of each active site, we carried out a “hand-over” assay, in which histidine-tagged Nsp12 used to NMPylate Nsp9 was subsequently removed, and another, untagged Nsp12 was added *post facto* (**Fig. 3C**). We found that the WT and D760A Nsp12s mediated de-NMPylation, whereas the D218A enzyme did not (**Fig. 3C**), ruling out an essential contribution of AS1 to the reversal of Nsp9 modification. Interestingly, while D760A is more efficient in NMPylating Nsp9 (**Fig. 1C**), it was slightly less efficient in the reverse direction. Thus, we cannot preclude te possibility of some involvement of AS1 in de-NMPylation, but the difference between the WT and D760A was barely significant (p=0.16), necessitating a more detailed analysis with additional variants of AS1 and AS2 residues.

Only a few examples of de-AMPylation are known, and most utilize different catalytic domains, either in the same or in different proteins (26). An example in which the same Fic domain mediates AMPylation and de-AMPylation of BiP, a major ER chaperone required for protein homeostasis in metazoans, has been recently reported (46). However, de-AMPylation releases AMP, not ATP, showing that FiD active site has both AMP transferase and phosphodiesterase activities (46). Future experiments will be required to reveal the mechanisms of reactions catalyzed by the NIRAN domain.

### Bisphosphonates inhibit Nsp9 modification

Strong inhibition of NMPylation reaction by PP_i_ (**Fig. 3B**) suggests that similar ligands that bind to AS2 (**Fig. 4A**) may competitively inhibit Nsp9 modification. To evaluate this possibility, we used chemically stable PP_i_ analogs bisphosphonates. We chose two FDA-approved compounds, Foscarnet (Fos) and Risedronate (Ris), as representative non-nitrogenous and nitrogenous bisphosphonates, respectively. Fos inhibits viral DNA polymerases, including HIV reverse transcriptase (47,48), and is used for treatment of infections caused by viruses in *Herpesviridae*. Ris is broadly used to treat diseases associated with bone loss, such as osteoporosis (49).

**Figure 4.**
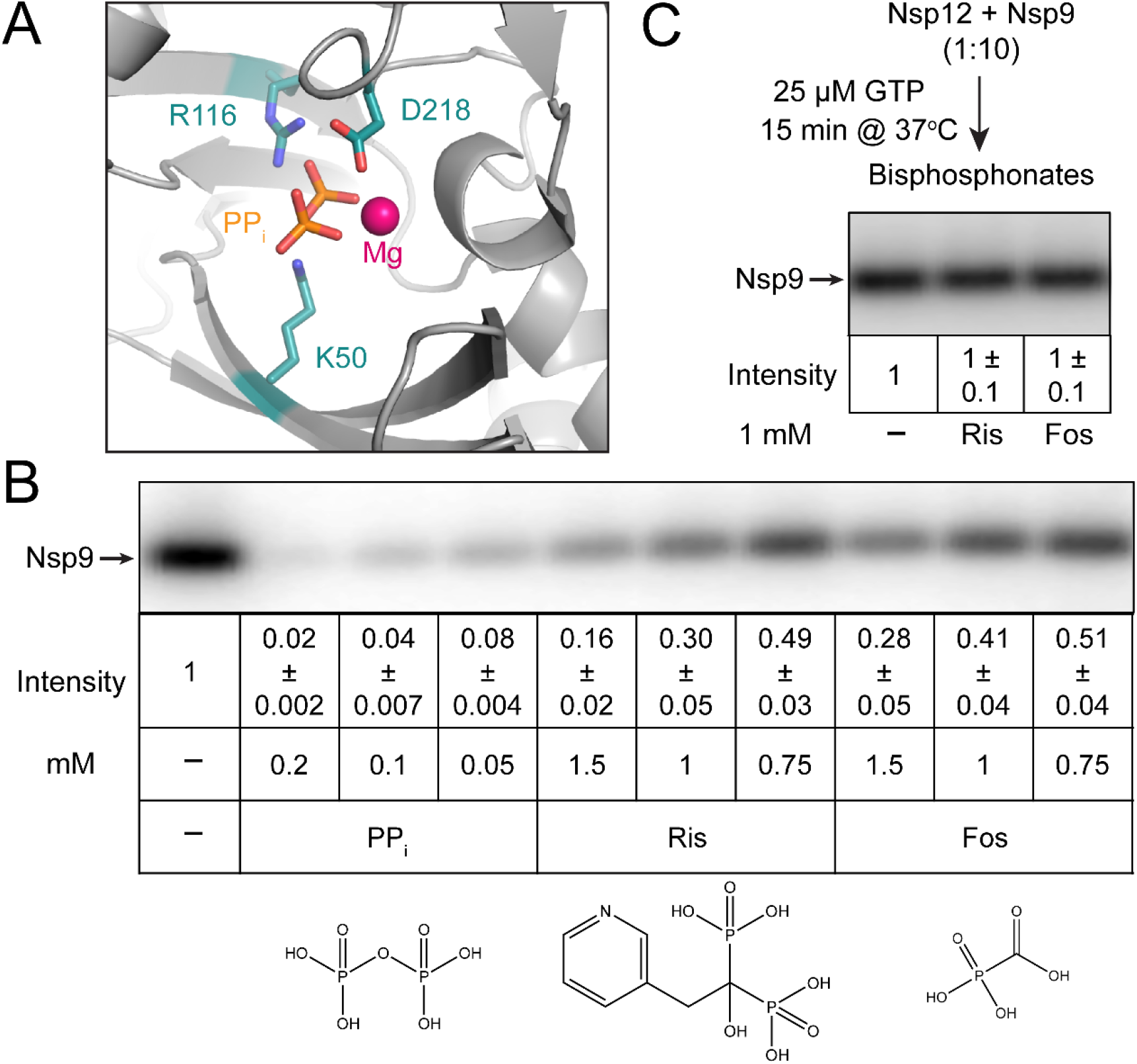
Effects of bisphosphonates on Nsp9 modification. **A**. Cryo-EM structure of the SARS-CoV-2 RdRp with PP_i_ bound to the NiRAN domain AS2. PDB: 7AAP (36). **B**. Inhibition of NMPylation by PP_i_ and bisphosphonates; structures are shown below each compound. **C**. Bisphosphonates do not promote the removal of NMP from modified Nsp9. In B and C, signal intensity was compared to that observed in the absence of added ligands (set at 1) and is shown as mean ± SD (n=3).

We show that Fos and Ris inhibit Nsp9 NMPylation, although less efficiently than PP_i,_ (**Fig. 4B**). While PP_i_ reduced NMPylation more than ten-fold when present at 50 μM, only two-fold inhibition was achieved at 0.75 mM of either bisphosphonate (**Fig. 4B**). These results are not surprising, given that PP_i_ reverses the reaction whereas bisphosphonates are expected to work merely as competitive inhibitors of the forward reaction. Indeed, unlike PP_i_, neither compound induced the removal of the GMP moiety from Nsp9 (**Fig. 4C**). We propose that bisphosphonates could be explored as inhibitors of NiRAN-mediated NMPylation; while neither of the two compounds tested was a potent inhibitor, many bisphosphonates are available or can be made to support structure-guided drug discovery.

## DISCUSSION

### Roles and targets of vital NiRAN NMPylation

The NiRAN domain is essential for replication of several human respiratory viruses, including the alphacoronavirus HCoV-229E, which causes the common cold, and betacoronaviruses SARS-CoV (18) and SARS-CoV-2 (19). Nucleotidylation activity of the NiRAN domain, which lacks any sequence homologs, was initially suggested by elegant bioinformatics analysis and confirmed by a proof-of-principle demonstration that EAV RdRp was capable of self-NMPylation (16–18). Structural similarities between SelO and NiRAN (16,17) further strengthened by identification of Nsp9 as a NiRAN target among HCoV-229E proteins (19), argue that the NiRAN domain is a protein NMPylase.

In their pioneering study suggesting and confirming the existence of the NiRAN domain, Lehmann et al. postulated three potential roles of NiRAN-mediated NMPylation in the nidoviral replicative cycle (18). One possible role is that of an RNA ligase, although the identity of the substrates, and indeed the step itself, remains entirely hypothetical to date. Another is that of a guanylyltransferase (GTase) involved in “capping” the 5′-end of transcribed RNA. Such capping is essential for viral replication and successful host infection, and all enzymes involved in the capping pathway, save the GTase, had already been identified years previously. The third possibility is that it serves a protein “primer” of RNA synthesis, by covalently binding a nucleotide and, following its extension to a dinucleotide, delivering it to the 3′-end of the viral RNA template. Such priming is widely used across viral families (50).

In discussing each of these putative roles, the authors noted that the sum total of structural, functional, and phylogenetic evidence then available, including their own findings, could not be entirely reconciled with any single role, much less definitively preclude the two others. Several recent studies have been less hesitant, assigning to NiRAN exactly one of these roles.

*First*, Yan *et al*. posited that NiRAN performs a “capping” role. In support of this assignment, they cited primarily structural arguments based on a cryo-EM snapshot of an extended RTC in which the N-terminus of Nsp9 was observed deep within the NiRAN active site, where it contacted a bound GDP molecule in a conformation stabilized by base-stacking with the His75 residue (20). They reasoned therefore that Nsp9 must be either the target of NiRAN NMPylation or a competitive inhibitor of it, concluding the latter since their functional assays detected the formation of capped RNA but not the NMPylation of Nsp9.

*Second*, Slanina *et al*. posited instead that NiRAN NMPylates Nsp9, which then serves as a primer of RNA synthesis (19). Their functional studies provided direct evidence of NMPylation of Nsp9 and NiRAN mediation thereof, with mutational and phylogenetic data supporting the additional conclusions that this NMPylation requires a free N terminus and allows little variation within the N-terminal tripeptide (**Fig. 1B**). In particular, the indispensability both of Asn2 for Nsp9 NMPylation *in vitro* and of NiRAN activity for viral replication provided a profound and elegant explanation why Asn2 is the only invariant residue across all Nsp9 homologs (19).

### Passing the baton: a speculative but integrative model

How can such findings be reconciled with one another, let alone with preceding or succeeding findings, including our own? Our results unequivocally demonstrate the importance of the native N-terminus of Nsp9 for its NMPylation (**Fig. 1C**), and thus we concur with Slanina *et al*. in arguing that the failure by Yan *et al*. to observe any such NMPylation is entirely due to their use of an artificial Nsp9. The conclusion put forward by the latter – that NiRAN must therefore cap 5′ pRNA, and do so directly – is thus unfounded. However, if NiRAN is not the GTase “missing link” in the capping pathway, no obvious candidate for this essential function remains.

As expected due to the lack of sequence-specific contacts between the nucleotide base and NiRAN residues (17,20), we found that all natural NTPs compete with GTP (**Fig. 2B**), suggesting that Nsp9 can be modified by any nucleotide, and their respective cellular abundances will largely determine the identity of the adduct. However, it is possible that AS2 specificity may be “tuned” in the presence of other RTC components.

We also show that, unlike the RNA chain synthesis, Nsp9 modification is readily reversible in the presence of PPi (**Fig. 4B**) and that Nsp9 interactions with AS2 are highly dynamic, i.e., NMPylated Nsp9 released from Nsp12 can be handed over to another enzyme for de-NMPylation (**Fig. 3C**). Finally, we show that ligands that bind to AS2, including nucleoside mono- and di-phosphates (**Fig. 2C**) and bisphosphonates (**Fig. 4B**), inhibit Nsp9 NMPylation.

Taken together, these results strongly argue for NMPylation of Nsp9 at NiRAN AS2. If so, to what end? Nsp9 binds RNA, with no apparent sequence specificity (27,29), and Nsp12 (20), but it is not clear how Asn1 modification would affect either interaction: residues thought to bind RNA are far away from Asn1 (27,29), and our results are inconsistent with any significant thermodynamic contribution of the NMPylation of Nsp9 to its binding to Nsp12 (**Fig. 1D**). Rather, Nsp9 appears to be ideally suited to deliver NMP to secondary acceptors: the NMP moiety is attached to the primary amine of N-terminal Asn1 (19) located at the end of a flexible N-terminal tail, and protein-N-NMP linkages are common in nucleotidyl transferases that catalyze ligation and capping reactions (51,52).

Therefore, we envision an essential role for NMPylated Nsp9 in *both* priming and capping (**Fig. 5**), perhaps as vital to the outcome as a baton passed between runners in a race. *First*, Nsp9 is NMPylated by the NiRAN domain at AS2 and then dissociates from Nsp12. *Second*, NMP-Nsp9 binds to AS1 and serves as a primer for RNA synthesis; although Nsp9 is not known to bind to specific RNA sequences (27,29), it is possible that, when bound to an RTC, NMP-Nsp9 recognizes a specific sequence/structure in the viral RNA to direct precise initiation. It is likely that different RdRp complexes synthesize (+) and (−) RNA strands, complicating this analysis. *Third*, as the nascent pRNA chain grows and is displaced from tRNA, pRNA-Nsp9 rebinds to AS2 and a second nucleotidyl transfer reaction takes place to cap the pRNA, releasing the unmodified Nsp9 and resetting the cycle.

**Figure 5.**
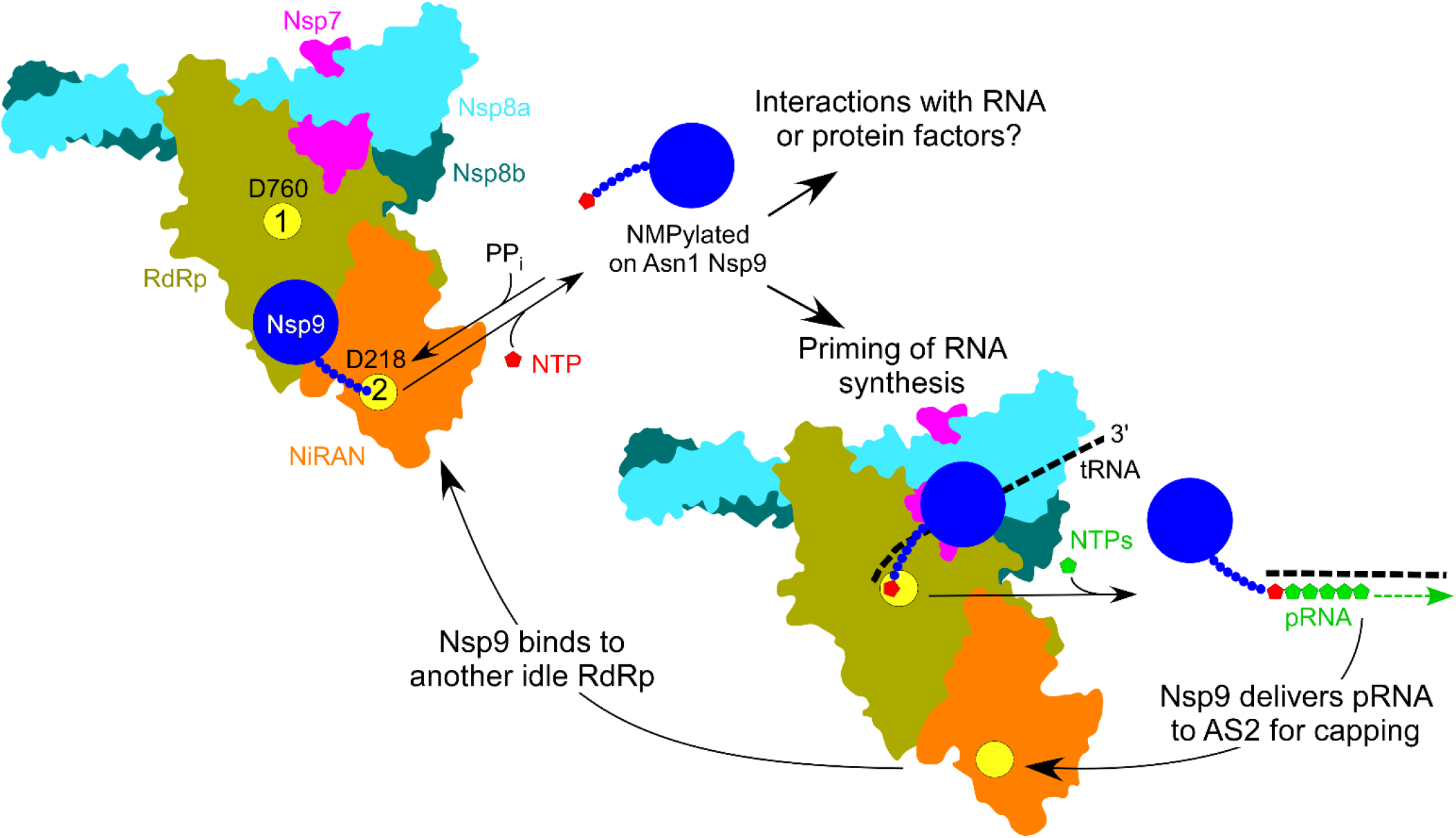
A model for Nsp9 cycle; see text for details. AS1 and AS2 are shown as yellow circles, with the key catalytic residues indicated.

### Avenues and implications for future research

We admit that this is a very speculative model and propose it to provoke investigation rather than to provide concrete answers. Mechanistic studies of SARS-CoV-2 RTCs are in their infancy, and future experiments will be needed to elucidate various aspects of its function and regulation. However, we argue that this model is a worthy starting point for several avenues of future research. Below we give several reasons for this claim and answer some anticipated objections.

*First,* such a capping mechanism is not unprecedented, for an analogous one has been described in rhabroviruses, such as vesicular stomatitis virus (VSV). VSV encodes a giant 2,100-residue L protein, which contains RdRp, nucleotidyl transferase, and methyl transferase modules (50). Via a covalent (L-histidyl-N^ɛ2^)-pRNA intermediate, L transfers the pRNA moiety to GDP to yield GpppA-RNA (50). Can NiRAN use GDP as an acceptor? We show that GDP competes with GTP during Nsp9 NMPylation (**Fig. 2C**) and concentration of GDP in infected cells may be sufficient (53). While very little is known about the NiRAN catalytic mechanism, other AMPylases possess surprising catalytic diversity: in addition to NMP, Fic proteins can transfer phosphocholine and phosphate to their targets (54,55).

*Second*, Nsp9 can be more than just a passive delivery vehicle for pRNA. Capping enzymes are composed of a nucleotidyl transfer domain fused to a distal OB-fold domain (51), suggesting that Nsp9 OB-fold domain (27,29), may cooperate with the NiRAN domain during pRNA capping, remodeling AS2. For example, Gre and TFIIS transcription factors, which reactivate arrested RNA polymerases in all domains of life, deliver the second catalytic Mg^2+^ ion to the active site to switch it into an RNA cleavage mode (44). Such remodeling of AS2, possibly in partnership with other components of the RTC, might mediate sequence specificity of NMPylation of Nsp9 in order to to prime the initiation of positive- versus negative-polarity RNA.

*Third*, we also admit that neither we nor others have definitively precluded all other possible protein targets of NiRAN. Enzymes that catalyze post-translational protein modifications, including AMPylation, have broad specificities – SelO was found to transfer biotin-AMP to a number of targets, including common control substrates, and only some cellular targets of SelO are thought to be genuine (21). In addition to Nsp9, NiRAN could also modify some other viral or host proteins, complicating the extension of *in vitro* results for an RTC to the viral replicative cycle or to the infection process as a whole. However, the high efficiency of NMP transfer to Nsp9 (**Fig. 1C**), conservation of Nsp9 N-terminus (**Fig. 1B**), and the essentiality of N-terminal Nsp9 residues for viral replication (19) all argue that Nsp9 is not only a true, but also perhaps the sole, protein target of NiRAN.

*Fourth*, our model has at minimum the virtue of not merely reconciling various seemingly contradicting findings, but also suggesting how they might be integrated into a more holistic understanding of the role of NiRAN, in concert with RNA and protein factors, in the entire nidoviral replicative cycle. We recently showed that over-optimization of the SARS-CoV-2 RdRp coding sequence to replace rare codons in a heterologous expression platform can lead to an inactive enzyme (22). We also showed that both AS1 and AS2 not only share substrates and inhibitors but also “cross-talk” via an allosteric pathway, and therefore that drug discovery and functional studies are myopic to focus exclusively on AS1 and unjustified in judging it to be the cause of all observed effects.

Similarly, a recent study found that whereas all existing cryo-EM structures of SARS-CoV-2 RdRp holoenzymes and RTCs modeled Nsp12 as chelating Zn centers, the physiological cofactors are in fact Fe-S clusters, which become replaced by Zn ions in the aerobic conditions in which proteins are typically purified (56). Furthermore, such clusters were found to be essential, for their disassembly via oxidative degradation inhibited both RdRp activity and viral replication. This result, obtained using a well-characterized nitroxide, suggests a potentially rich vein of COVID-19 therapeutics that might have been completely overlooked had the suitability of the cryo-EM structural preparations not been properly questioned.

Both these previous results and those presented here clearly demonstrate the inherent dangers in using reductionist approaches to draw conclusions for more complex and holistic systems, such as viral replicative cycles and processes of host infection. Such approaches have advantages for quickly yielding insights into a narrow and well-defined question, and so it is wholly understandable why they are particularly attractive for research into systems like SARS-CoV-2 RdRp, where pressing concerns motivate researchers to obtain practical results as rapidly as possible. On the other hand, such frenetic research can easily outpace the self-correction normally occurring in science, suggesting that wherever possible, researchers should strive to holistically validate reductionist findings (e.g. verifying replication of mutant viruses in cell culture) and clearly communicate aspects of research methods that might be expected to restrict the applicability of their results.

## SUPPLEMENTAL INFORMATION

Supplemental Information includes four figures and one table.

## ACKNOWLEDGEMENTS

We are grateful to Georgi Belogurov and Markus Wahl for helpful suggestions. This work was supported by a seed grant from The Ohio State University Office of Research and the National Institutes of Health grant GM067153 to I.A.

