## SUPPLEMENTARY INFORMATION for "NMPylation and de-NMPylation of SARS-CoV-2 Nsp9 by the NiRAN domain"

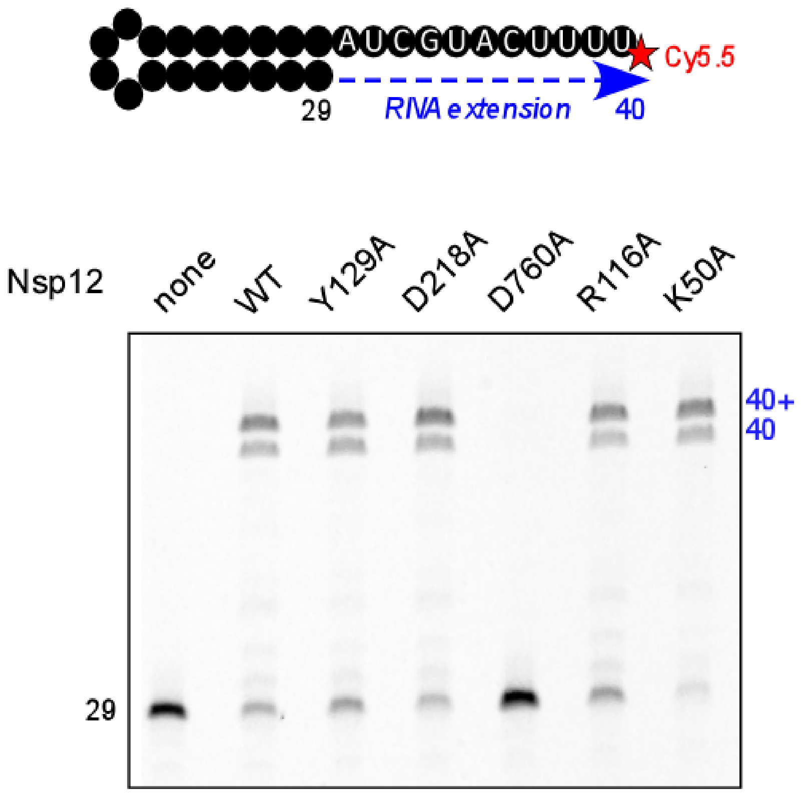


**Figure S1. RNA synthesis activities of Nsp12 variants.** RdRp holoenzymes Nsp7•$8_2$•12 (WT or indicated mutants) were assembled and tested as described previously (1). An 29-nt RNA hairpin scaffold that contains Cy5.5 at the 5' end is extended by RdRp to produce a 40-nt product; additional extension is thought to be mediated by Nsp8 after the completion of RNA synthesis (2).

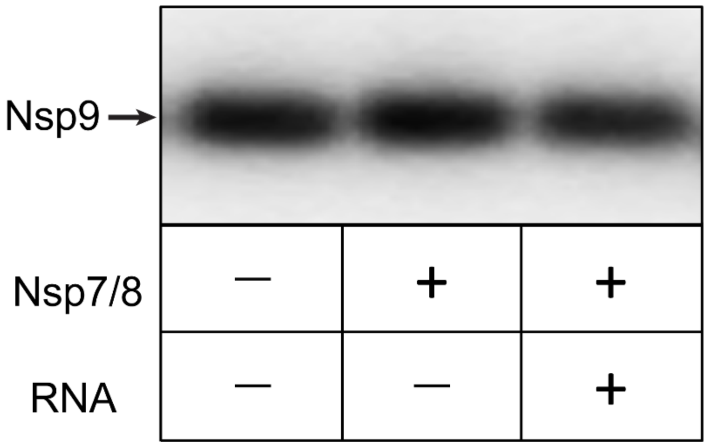


**Figure S2. Nsp12-mediated NMPylation of Nsp9 is observed in the holoenzyme and the transcription complex.** The NMPylation reaction was carried out by Nsp12 alone as in **Fig. 1** (first lane), in the presence of Nsp7 and Nsp8 added at 1.5 and 3x molar excess relative to Nsp12, respectively, to assemble the RdRp holoenzyme Nsp7•$8_2$•12 (second lane), and in a transcription complex assembled on the RNA scaffold shown in **Fig. S1** (third lane).

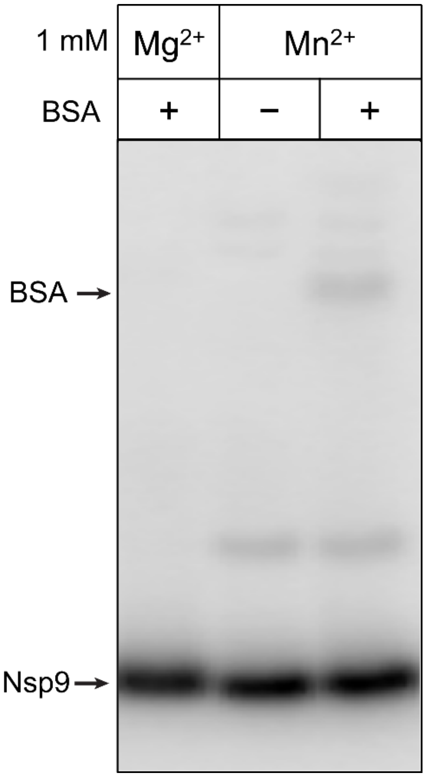


**Figure S3. Nsp12 mediates fortuitous NMPylation in the presence of the $Mn^{2+}$ ion**. We have not observed nucleotidyl transfer by Nsp12 to proteins other than Nsp9 in the presence of either 1 or 2 mM $Mg^{2+}$thought to represent physiological cellular levels (3); higher concentrations were not tested. However, we observed very inefficient NMPylation of Nsp12 (**Fig. 1C**) and BSA at 1 mM $Mn^{2+}$. 5 µM BSA, 0.5 µM Nsp12, and 5 µM Nsp9 were incubated for 10 min in the NMPylation buffer (see the standard NMPylation assay in the Method). An additional species visible on a gel is either Nsp9 dimer or uncleaved SUMO-Nsp9 fusion.

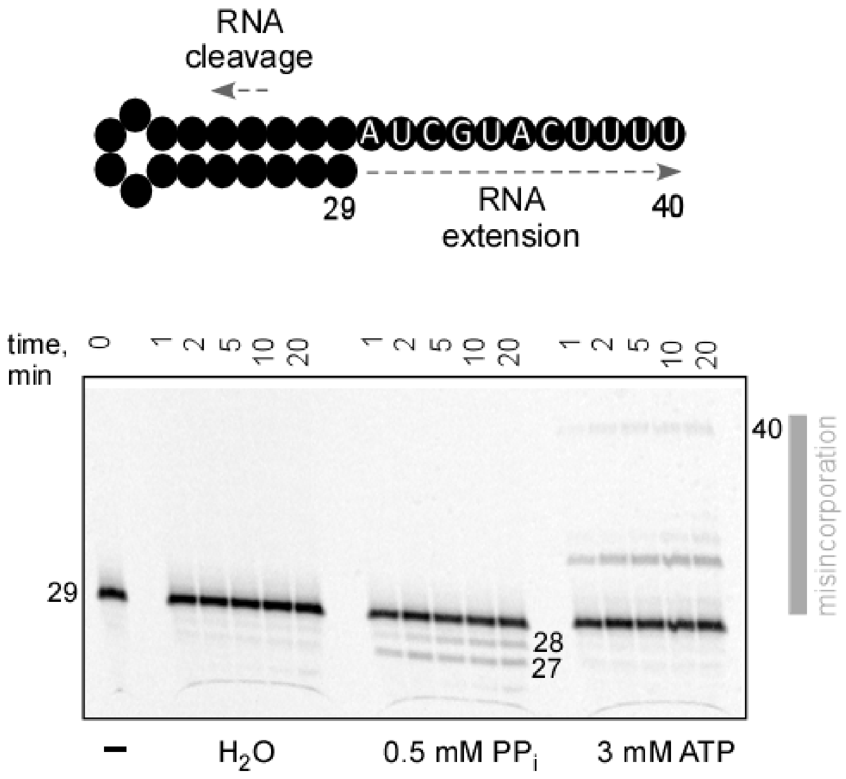


**Figure S4. SARS-CoV-2 RdRp does not mediate reverse pyrophosphorolysis in the presence of the noncognate substrate**. The reactions were carried out in the presence of 5 mM $Mg^{2+}$ to avoid titrating the metal ion by ATP. We observe synthesis of full-length 40-mer product, indicating that RdRp misincorporates AMP in place of UMP (1$^{st}$ position), GMP (3$^{rd}$ and 7$^{th}$ positions) and CMP (4$^{th}$ position) at high ATP concentrations.

**Table S1. Plasmids**

| Name | Key features/sequence | Source; Addgene# |
|---|---|---|
| pIA1362 | T7 promoter–$His_8$-GB1-TEV-Nsp7 | (1); 166860 |
| pIA1363 | T7 promoter–$His_8$-GB1-TEV-Nsp8 | (1); 166861 |
| pIA1364 | T7 promoter–$His_8$-GB1-TEV-Nsp9: expresses $^{GSN}$Nsp9 | This work; 172401 |
| pIA1400 | T7 promoter–$His_{10}$-SUMO-Nsp12 | This work; 172519 |
| pIA1402 | T7 promoter–$His_{10}$-SUMO-Nsp12[Y129A] | (1); 172403 |
| pIA1405 | T7 promoter–$His_{10}$-SUMO-Nsp12[D218A] | (1); 172404 |
| pIA1408 | T7 promoter–$His_{10}$-SUMO-Nsp12[D760A] | This work; 172405 |
| pIA1414 | T7 promoter–$His_{10}$-SUMO-Nsp9: expresses the native $^{N}$Nsp9 | This work; 172402 |
| pIA1420 | T7 promoter–$His_{10}$-SUMO-Nsp12[R116A] | This work; 172406 |
| pIA1421 | T7 promoter–$His_{10}$-SUMO-Nsp12[K50A] | This work; 172407 |
